# Rgg2/Rgg3 quorum sensing is a determinant of *Streptococcus pyogenes*-host interactions in a murine intact skin infection model

**DOI:** 10.1101/2023.07.14.549100

**Authors:** Reid V. Wilkening, Christophe Langouët-Astrié, Morgan Severn, Michael J Federle, Alexander R. Horswill

## Abstract

*Streptococcus pyogenes* is an obligate human pathobiont associated with many disease states. Here, we present a novel model of *S. pyogenes* infection using intact murine epithelium. From this model, we were able to perform RNA sequencing to evaluate the genetic changes undertaken by both the bacterium and host at 5- and 24-hours post infection. Analysis of these genomic data demonstrate that *S. pyogenes* undergoes significant genetic adaptation to successfully infect the murine epithelium, including changes to metabolism and activation of the Rgg2/Rgg3 quorum sensing (QS) system. Subsequent experiments demonstrate that an intact Rgg2/Rgg3 QS cascade is necessary to establish a stable superficial skin infection. Furthermore, activation of this pathway results in increased murine morbidity and increased bacterial burden on the skin. This phenotype is associated with gross changes to the murine skin, as well as histopathological evidence of inflammation. Taken together, these experiments offer a novel method to investigate *S. pyogenes*-epithelial interactions and demonstrate that a well-studied QS pathway is critical to a persistent infection.

**Importance:** *Streptococcus pyogenes* remains a pathogen of global importance, with significant total disease burden worldwide. Much of this burden is due to skin infection or sequalae thereof, yet little is currently known about the initial interactions between the organism and host skin. Here we present a new mouse model of skin infection. From this model, we were able to study gene expression by both the bacteria and the host during early infection time points. Both genetic and phenotypic data derived from these results demonstrate that a well-conserved *S. pyogenes* communication network, the Rgg2/Rgg3 quorum sensing pathway, contributes to establishing and maintaining a durable skin infection. We propose that by better understanding the genetic pathways needed to colonize and adapt to new niches, new approaches to preventing and treating *S. pyogenes* may be possible.

## Introduction

*Streptococcus pyogenes* (historically known as Group A *Streptococcus* or GAS) is an obligate human pathobiont associated with numerous disease states. *S. pyogenes* causes human illness ranging from superficial infections of the pharynx (e.g. Streptococcal pharyngitis) and skin (e.g. impetigo, cellulitis) to life-threatening invasive or toxin-mediated manifestations (e.g. necrotizing fasciitis, streptococcal toxic shock syndrome, scarlet fever and septic shock). *S. pyogenes* is also the causative agent of clinically relevant non-suppurative sequalae, including post-streptococcal glomerulonephritis and rheumatic heart disease [1, 2]. Historically, streptococcal research efforts have primarily focused on understanding the molecular mechanisms of invasive illness, unsurprising given the severity of these diseases. Despite the fields’ focus on invasive illness, it is estimated that 5-20% of school age children carry *S. pyogenes* asymptomatically in their nasopharynx [3, 4]. Less is known regarding skin carriage and infection rates. Large microbiome studies typically have not had the resolution needed to delineate *S. pyogenes* from other Streptococci or Firmicutes, leaving a considerable knowledge gap. Other studies have also demonstrated that skin microbiome shifts with age, with Firmicutes more common during childhood, though the reasons for this transition remain unclear [5, 6]. Nonetheless, years of studies demonstrate that *S. pyogenes* can be endemically carried on the skin, with carriage contributing to both superficial infection and high rates of non-suppurative sequelae.

Despite the foundation established by descriptive studies, there is little known about the genetic determinants that enable *S. pyogenes* to colonize the skin. It is proposed that the transition from colonizer to pathogen may be mediated by natural breaks in the skin (such as insect bites, minor trauma, or scabies infections), though mechanisms remain under investigation [7]. For example, it has been postulated that scabies and streptococcal coinfections are frequently seen due to complement inhibition by the scabies mite [8]. Despite our extensive knowledge regarding the genetic determinants of invasive *S. pyogenes* illness using models of necrotizing fasciitis or other models bypassing the integument [9, 10], less is known about bacterial infection of unbroken skin.

In this study, we present a model of *S. pyogenes* infection of intact murine epithelium. From this model, we performed dual RNA sequencing (RNAseq) to study transcriptional changes occurring during interaction between the bacteria and the murine host. Our data demonstrate that the Rgg2/Rgg3 quorum sensing (QS) system is the most upregulated pathway in *S. pyogenes* during adaptation to skin. *In vivo* experiments using *S. pyogenes* NZ131 with intact and abrogated QS functionality show that the presence of an intact QS cascade led to a more protracted duration of carriage with greater bacterial burden, and increased morbidity. These findings have important implications for both our understanding of *S. pyogenes* carriage as well as novel treatment options to counter skin infections.

## Materials and Methods

### General conditions for cultures

Bacterial strains used during these experiments are described in Table 1. Strains were grown under typical laboratory conditions. All GAS strains were grown in Todd Hewitt medium (Research Products International) supplemented with yeast extract (Todd Hewitt Yeast, THY). Broth cultures were grown statically at 37°C. When needed, THY plates were made by adding agar to final concentration of 1.4%. CHROMagar StrepA medium (CHROMagar) was used to isolate GAS from mouse infections. Plates were allowed to incubate at 37°C + 5% CO2 for approximately 24 hours following inoculation to allow for full chromogenic development. When appropriate, antibiotics were supplemented at the following final concentrations: chloramphenicol 10 μg/ml, erythromycin, 1 μg/ml.

**Table 1.** Strain used in this study.

| Strain | Description | Reference |
| --- | --- | --- |
| NZ131 | Wild-type M49 *S. pyogenes* isolate | [19, 52] |
| Δrgg3 | NZ131 Δrgg3::cat | [25] |
| Δrgg3 shp2<sup>GGG</sup>shp3<sup>GGG</sup> | NZ131 Δrgg3::cat, shp2<sup>ATG@GGG</sup>, shp3<sup>ATG@GGG</sup> | [26, 53] |
| NZ131 P<sub>shp3</sub>-luxABCDE | NZ131, P<sub>shp3</sub>-luxABCDE integrated at T12 phage site attB | [18] |

### Animal work and ethics statement

Mice were housed in the BSL2 facilities provided by the University of Colorado School of Medicine. All experiments were conducted under approved institutional animal care and use protocol #00941. Food and water were provided *ad libatum* during experiments, with supplementary dry feed provided on the floor of cages during extended infection experiments. At end points, mice were euthanized via CO2 inhalation with cervical dislocation as a secondary method. Male 7-week-old C57BL/6 mice obtained from Jackson Laboratory were used for all experiments. Mice were allowed to acclimate for at least one week in the vivarium prior to experimentation. One day prior to inoculation, the dorsum of the mice was shaved using electric hair trimmers. Residual hair was removed with Nair (Church and Dwight Co., Inc.) and mice were allowed to recover overnight before inoculation.

### RNA preparation and sequencing

To prepare samples for RNA sequencing, NZ131 cultures were grown as noted above. Independent cultures were prepared to represent 3 biological replicates and were processed in parallel. Input control samples were grown to mid-log in THY broth with an optical density (OD_600_) ranging from 0.4-0.5. At this time, samples were removed and preserved with RNAprotect (Qiagen) per manufacturer’s instructions. All culture densities were determined by serial dilution and plating on THY agar plates. Cultures used for *in vivo* infection were grown in a similar manner, then prepared for inoculation via centrifugation and resuspension in PBS to a final concentration of ∼5×10^8^ CFU/ml. A final volume of 200 μl of culture was pipetted onto the pad of sterile bandages, which were affixed to mice overlying depilated areas and secured with a second bandage. All bandages were left in place for 5 hours prior to removal. After 5 or 24 hours of recovery post bandage removal, mice were euthanized with CO_2_. Their skin was swabbed 10 times with a sterile cotton swab pre-wetted in PBS to quantify bacterial burden then a 6 mm punch biopsy was used to isolate tissue samples. These samples were stored in RNAprotect at -80°C prior to processing. To obtain sufficient bacterial RNA, skin samples from 5 mice were pooled to generate a single sample. To extract RNA, skin samples were homogenized with 1 mm zirconia-silicate beads. After removing insoluble material by centrifugation, the resulting supernatant was then applied to 0.1 mm zirconia-silicate beads and mixed with RLT buffer (Qiagen). RNA isolation proceeded as per manufacturer’s instructions using a RNeasy Plus kit (Qiagen). Samples were DNAse I treated, and RNA was quantified on a Nanodrop One spectrophotometer prior to submission to the Microbial Genome Sequencing Center (MiGS, now known as SeqCenter, Pittsburgh PA). MiGS undertook ribosomal RNA depletion via RiboZero Plus depletion (Illumina), Stranded Total RNA Prep Ligation with IDT for Illumina Indices library preparation and paired sequencing on an NextSeq2000 platform. Post sequencing, bcl2fastq (Illumina, v2.20.0.422) was used to demultiplex and trim adapters, and to quantify reads. Bacteria-only inputs underwent a minimum of 12 million reads per sample, while samples containing both murine and bacterial RNA underwent a minimum of 50 million reads per sample.

### Bioinformatics

Transcriptomic results were processed using CLC Genomics Workbench (Qiagen, v20.0.04). Raw unpaired data were imported, adapter trimmed, aligned, paired, and annotated into CLC Workbench using default settings: mismatch cost, 2; insertion and deletion cost, 3; length and similarity fraction, 0.8. Reads were annotated to reference genomes in a sequential fashion. First, all reads were aligned to the *Mus musculus* genome (mm10). Unmapped reads to the *M. musculus* genome were then aligned to the *S pyogenes* NZ131 genome (NC_011375.1). Uniquely mapped mouse or bacterial transcripts were normalized and used for differential gene expression analysis using R (v4.2.2), Rstudio (v2022.12.0+353, RRID: SCR_000432), and DESeq2 (v1.36.0, RRID: SCR_015687) ([11, 12]. Differentially expressed genes were filtered based on an absolute log2 fold change cutoff and Benjamini-Hochberg False Discovery Rate (FDR) cutoff. fGSEA (v1.22.0, RRID: SCR_020938) with 10,000 permutations and the Hallmarks and GO Biological Processes gene set collections from the Molecular Signatures Database were used for mouse pathway analysis [13–15]. NZ131 pathway analysis was done by Kegg pathway analysis [16, 17]. Mouse and bacterial RNA sequencing data, including raw sequencing files and post-analysis results, were deposited to National Center for Biotechnology Information Gene Expression Omnibus (GEO) with accession number GSE237018.

### *In vivo* imaging experiments

To study *in vivo* quorum sensing activity, we utilized the P_shp3_-luxABCDE plasmid (pRVW14) reporter strain first described [18] in the WT NZ131 background. Input strains were grown to mid-log prior to back-dilution in PBS to ∼3.5×10^8^ CFU/ml. Mice were anesthetized with isoflurane. 300 μL of bacteria were applied to sterile bandages and applied to depilated mice. Bandages were then covered with a Tegaderm occlusive dressing (3M). Mice were imaged hourly at 0-4 and 24 hours post inoculation using a Xenogen IVIS-200 (Perkin-Elmer) imaging system. Mice were allowed to recover from anesthesia between time points. Data analysis was conducted using the Living Image software package and all flux data are reported in photons/second/cm^2^/steradian (p/s/cm²/sr).

### Longitudinal Mouse Colonization Experiments

Experiments to assess the impact of quorum sensing on murine colonization were conducted as described above with the following modifications. As described previously, input bacterial cultures were grown to mid-log phase and diluted to attain approximately 1×10^8^ CFU/200 μL inoculum. Samples were applied to sterile bandage pads and affixed to depilated mice. Bandages were then covered with a Tegaderm occlusive dressing (3M) and secured with a second bandage. At each time point, this dressing was removed, mice were weighed, swabbed with a pre-wetted cotton swab, and a new dressing was affixed. Swabs were placed in sterile PBS, vortexed for 30 seconds each, serially diluted and plated on selective *S. pyogenes* CHROMagar to assess bacterial burden. At each time point, representative photos of the mice were taken.

On the final day of the longitudinal colonization experiment, following swabbing 1 mouse/cage (3 mice/group) was randomly selected for histological analysis. Skin from the dorsum of the mouse was excised. Skin was placed in histology cassettes and immediately fixed in 4% paraformaldehyde for a minimum of 48 hours prior to storage in 70% ethanol. Tissue samples were processed by the Histology Core at the University of Colorado. Samples were embedded in paraffin and sectioned. Samples were subjected to hematoxylin and eosin staining. Adjacent sections were deparaffinized in citrate and stained with monoclonal mouse anti-F4/80 antibody at a 1:400 dilution (BioLegend), or polyclonal goat anti-MPO antibody at a 1:500 dilution (R&D Systems). Representative images were captured using a Keyence BZ-X700 microscope at indicated magnifications.

### Statistics

Statistical analyses were performed in Prism 9 (Graph Pad). The Kruskal-Wallis tests with Dunn’s correction for multiple hypotheses with an alpha threshold level of 0.05 was used to analyze CFU and weight data. IVIS luminescence data were analyzed with a two-way ANOVA test with Geisser- Greenhouse correction and Dunnett’s multiple comparison test.

## Results

### *Streptococcus pyogenes* can transiently colonize intact murine epithelium

We first set out to develop a model of *S. pyogenes* colonization of intact, uninjured murine epithelium, as we hypothesized such a model would be necessary to understand early host-bacterial interactions (Figure 1A). It was necessary to remove murine hair to allow direct interaction with the epithelium, but we otherwise sought to provide an unmodified substrate. We chose to work with the M49 strain NZ131 due to the genetic tractability of this strain, the availability of complete sequencing and the fact that serotype M49 strains have been shown to be generalists capable of infecting both skin and pharyngeal sites [19–21]. We utilized an indirect approach, whereby NZ131 was inoculated onto the sterile pad of a band-aid. This band-aid was then affixed to the depilated dorsum of mice and left in place for 5 hours prior to removal. Preliminary studies demonstrated that such an approach was able to maintain detectable levels of colonization for up to 24 hours (Figure 1B). When this model was carried out longitudinally beyond 24 hours, there was an approximate loss of colonization at 1-1.5 log per day (data not shown).

**Figure 1:**
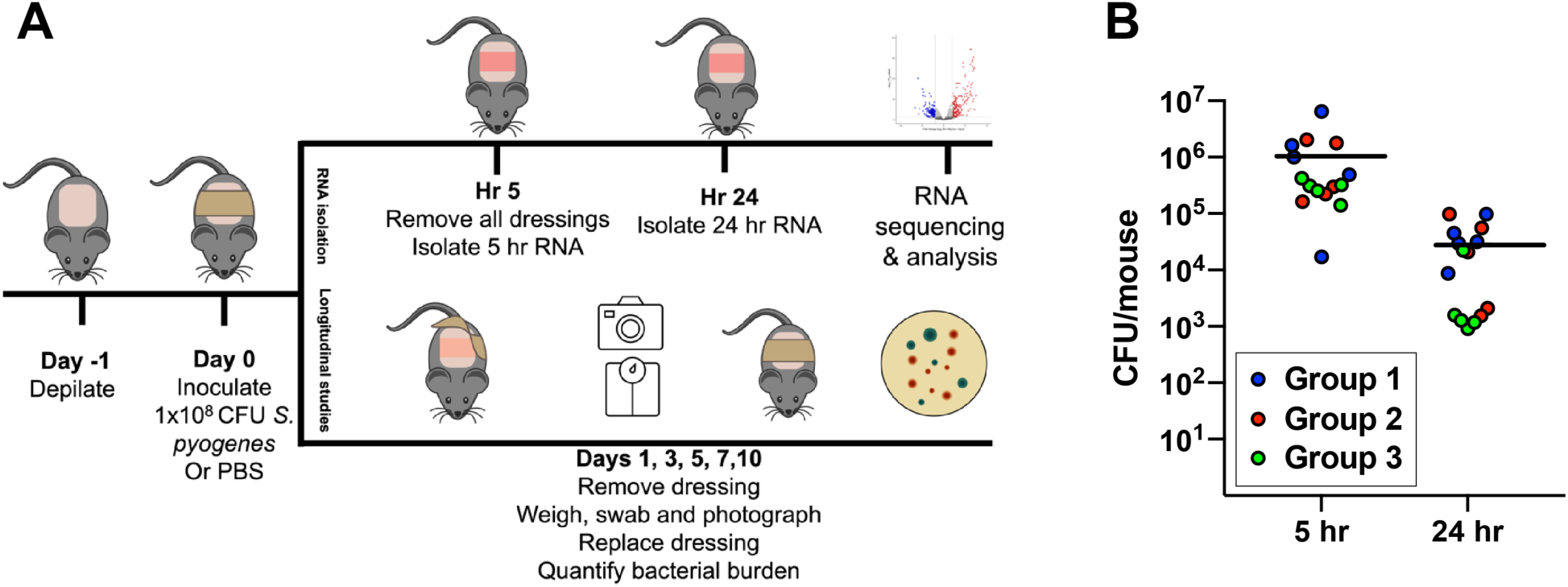
**A.** Experimental outline. All experiments begin with depilation on day -1 and inoculation on day 0. Upper limb demonstrates approach used for RNA sequencing experiments. Lower limb demonstrates approach used for longitudinal experiments. **B.** CFU per mouse at 5 and 24 hours, at time of RNAseq sample collection. Each group represents 5 mice, pooled for RNAseq to generate 3 biological replicates. Black bar represents mean of all 15 mice for a given timepoint.

### Dual RNAseq of NZ131 murine skin colonization

We next sought to collect both murine and bacterial RNA for transcriptomic analysis at the 5- and 24-hour time points. Using the above-described infection model, mice inoculated with wild type (WT) NZ131 were sacrificed at 5 and 24 hours after inoculation and RNA from the murine epithelium and the associated bacteria was extracted. Samples from mice mock infected with PBS were also obtained at each time point, as were pre-inoculum NZ131 input controls. Raw sequencing data was sequentially aligned, first to the *Mus musculus* genome, then remaining transcripts aligned to the NZ131 genome. Analysis of expression data were conducted as described above and completed analyses are presented in tables S1 and S2.

### NZ131 rewires metabolic pathways during murine skin adaptation

Principal-component analysis (PCA) of bacterial data demonstrates that *S. pyogenes* undergoing murine adaptation are more similar at 5- and 24-hour samples than their cognate input control, as demonstrated by the 5- and 24- hour samples uniquely clustering from the input cultures. (Figure 2A). Using an FDR threshold P-value of ≦0.05 and a log_2_ fold change 2|2|, there were 166 genes upregulated and 126 genes downregulated at 5 hours versus input control, denoting significant change in approximately 16% of the genome. At 24 hours, there were 159 genes upregulated and 88 genes downregulated versus input control, significant change in around 14% of the total genome (Figure 2B/C, Table S3/S4). There was notable overlap between differentially expressed genes (DEGs) across both time points (Figure 2D). Between pooled biological replicates, there was good concordance in differential gene expression across replicates at 5 and 24 hours (Figure 2E/F)

**Figure 2.**
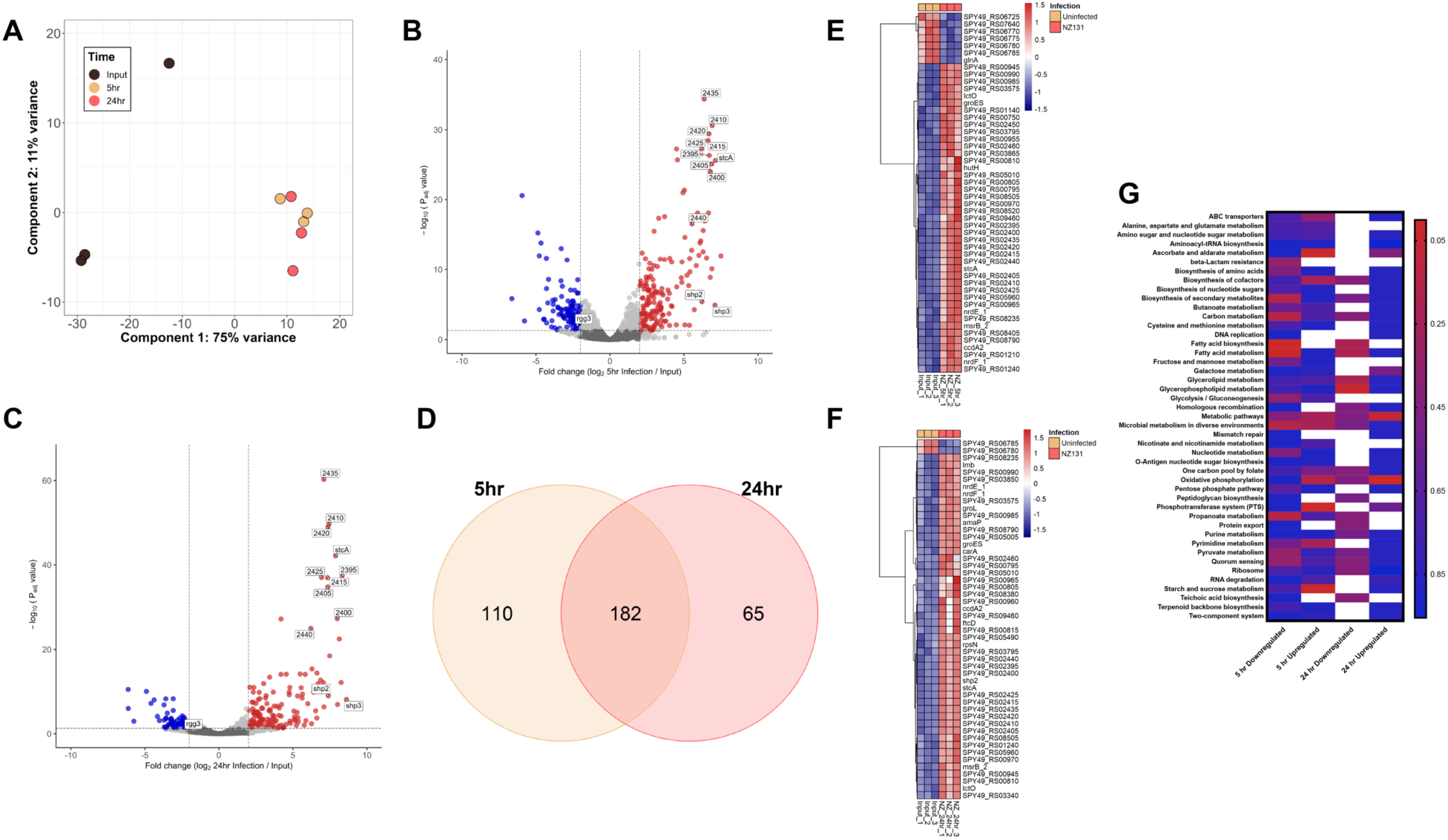
**A:** PCA analysis of NZ131 grown in THY broth (black) and after 5 hours (gold) and 24 hours (pink) of murine skin infection. **B, C:** Volcano plots demonstrating the distribution of differentially regulated genes at 5- and 24-hours post skin infection. Genes that are part of the Rgg2/Rgg3 quorum sensing pathway are labeled with locus names. **D:** Venn diagram demonstrating differentially regulated genes at 5 versus 24 hours post inoculation onto murine skin. **E, F:** Heatmap of the top differentially expressed *S. pyogenes* genes which are normalized counts transformed with Z-score method. Each column represents a biologically unique sample, with input controls on the left and 5 or 24 hour samples on the right. **G:** KEGG pathway analysis of NZ131 gene expression at 5- and 24-hours post inoculation versus input control of the top 45 regulated pathways based on log_2_ FC>|2|.

To begin understanding organism-level genomic adaptations occurring during murine skin colonization, DEGs were subjected to pathway enrichment analysis against the Kyoto Encyclopedia of Gens and Genomes (KEGG) [16, 17, 22]. Using an FDR P-value ≤0.05, there were 3 and 2 pathways upregulated at 5 and 24 hours respectively, and 2 and 1 downregulated at 5 and 24 hours respectively (Figure 2G). KEGG analysis of the upregulated genes at 5 hours demonstrated significant enrichment of ascorbate and aldarate metabolism, starch and sucrose metabolism and the phosphotransferase system. At 5 hours, there was significant downregulation of pathways associated with fatty acid biosynthesis and metabolism. At 24 hours, oxidative phosphorylation and metabolic pathways were upregulated, while glycerophospholipid metabolism were downregulated. These results demonstrate that *S. pyogenes* is subjected to metabolic stress during adaptation to growth on murine epithelium, decreasing metabolic use of fatty acids, while increasing utilization of non-glucose carbon sources. These results were similar with the differentially expressed pathways described in Cook *et al*., which used a vaginal colonization model of NZ131 infection and found a decrease in fatty acid biosynthesis gene expression and an increase in ascorbate metabolism genes expression [23]. This suggests that despite the differences between mucosal vaginal epithelium versus keratinized epithelium, available nutrients and stressors are similar enough to require significant metabolic adaptation on the part of *S. pyogenes* to tolerate these environments.

As we had envisioned our model as one of adaptation to uninjured epithelium, we next investigated whether canonical *S. pyogenes* virulence factors were induced during our study period. Review of RNAseq data do not show a marked, consistent expression increase in the majority of virulence-associated genes. *slo* (Streptolysin O)*, scpC* (SpyCEP) and *sdaB* (Streptodornase) are modestly upregulated at 5 and 24 hours (2-3 fold). *nga* (NAD glycohydrolase) is upregulated at 5 but not 24 hours. *sls* (Streptolysin S)*, ska* (Streptokinase)*, enn* (M-related protein)*, speB* (cysteine protease) and the *has* (hyaluronic acid) operon do not show significant changes in expression between broth culture and murine isolates at any time point (Table S3/S4). Other genes known to be regulated by the *covRS* operon, such as *pepO* (Endopeptidase PepO)*, arcA* (Arginine deaminase) and *sclA* (cell surface anchored collagen-like adhesin) do not show significant upregulation at either 5 or 24 hours [24]. Taken in whole, these results suggest that *S. pyogenes* virulence factors do not play a central role in early murine skin adaptation.

### The Rgg2/Rgg3 quorum sensing pathway is up-regulated during adaptation to murine skin

At both 5- and 24-hour time points, the Rgg2/Rgg3 quorum sensing pathway was markedly upregulated. This pathway is well-conserved across all sequenced *S. pyogenes* isolates and is well described in [18, 25–30]. **In** brief, the QS cascade uses two peptide pheromones, Shp2 and Shp3, to interact with a pair of transcriptional regulators, Rgg2, a positive regulator and Rgg3, a negative regulator (Figure 3). Phenotypically, these operons are now known to increase lysozyme resistance (mediated by *stcA*), alter immune regulation (mediated by the *qim* operon downstream of *shp3*), decrease *slo* expression (mediated by *qimH*) and improve NALT colonization. All the genes directly regulated by this pathway, as well as the previously described phenotypes, are contained within the operons downstream of each of the peptide pheromones [18, 25, 31–33]. In our RNAseq data, we found that s*hp3* was upregulated 136-fold at 5 hours and 392-fold at 24 hours. Similarly, *shp2* was upregulated 75-fold at 5 hours and 169-fold at 24 hours (Table 2). Genes in the associated operons downstream of the pheromones were also significantly upregulated (Figure 3, Table 2). This induction does not appear to be due to an increase in the positive regulator of QS *rgg2,* which was not differentially regulated versus input at either 5 or 24 hours. However, the negative transcriptional regulator of QS, *rgg3,* was significantly downregulated approximately 5-fold at both time points, suggesting a possible mechanism by which skin colonization leads to Rgg2/3 induction.

**Figure 3:**
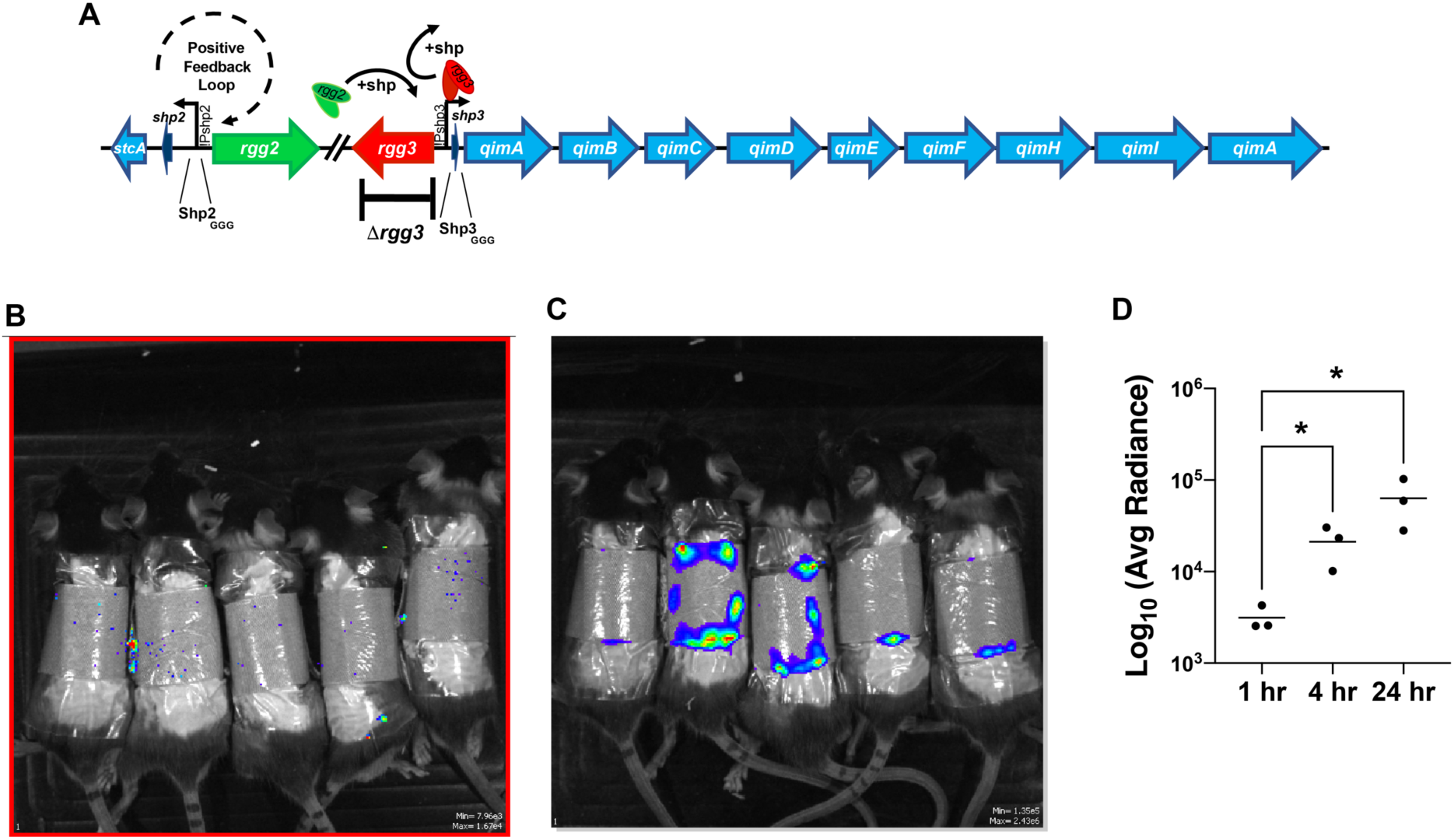
**A**. Schematic demonstrating the Rgg2/Rgg3 quorum sensing pathway and associated operons. Activation of the pathway leads to de-repression of promoters P_shp2_ and P_shp3_ by displacement of Rgg3 (red) by Rgg2 (green) upon Shp peptide binding. Pathway activation leads to significant upregulation of both of the Shp peptides (dark blue), perpetuating a positive feedback loop. There is also upregulation of associated downstream genes (light blue). Of note, *rgg2* was not differentially regulated in this dataset, though *rgg3* was found to be repressed 5-fold in mice as compared to broth culture. Mutations used in longitudinal infection experiments are highlighted below the operon. **B/C.** Representative IVIS images at 1- and 24-hours post inoculation. Mice were inoculated with NZ131 with integrated P_shp3_-*lux_ABCDE_* transcriptional reporter and monitored longitudinally via IVIS *in vivo* imaging. **D.** Rgg2/3 QS pathway is significantly induced during murine skin infection. Each point represents average radiance for 5 mice treated as technical replciates. Radiance reported as [p/s/cm²/sr]. * p<0.05.

**Table 2.**
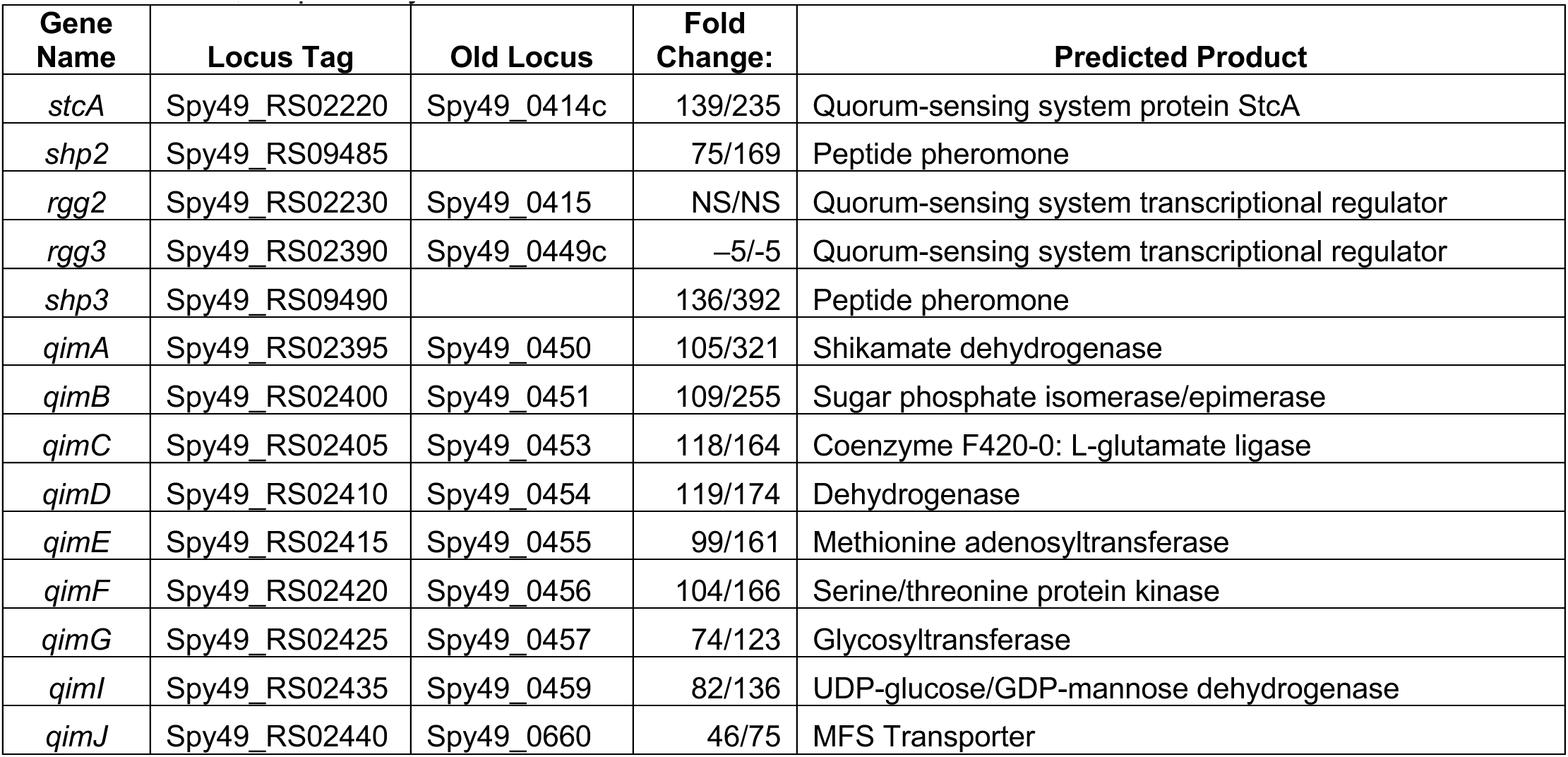
Genes associated with the Rgg2/Rgg3 regulon. Locus tag refers to current NCBI annotation. Old locus tag refers to original NCBI annotation. Fold change is shown at 5 hours and 24 hours, respectively.

To confirm the RNAseq results, we inoculated mice with WT NZ131 bearing a P_shp3_-lux transcriptional reporter and imaged mice serially using an IVIS system to interrogate activation of the QS cascade. As expected, no signal was produced initially. Induction was noted to begin around 4 hours, approximately consistent with the timing of induction observed in our RNAseq dataset. There was a statistically significant difference in QS activation at 24 hours when compared to 1-hour post-inoculation (Figure 3B,C). These data confirm our RNAseq and phenotypic results and demonstrate that murine-bacterial interactions activate Rgg2/3 QS at the transcriptional level.

### NZ131 skin infection stimulates an inflammatory response *in vivo*

As we performed dual RNAseq, we also investigated host responses within the murine epithelium to NZ131. PCA suggests that duration of infection leads to unique gene expression profiles (Figure 4A,B). Using the same thresholds as chosen previously, (FDR P-value ≤0.05 and a log_2_fold change ≥|2|), there were 39 DEGS at 5 hours and 76 DEGs at 24 hours (Figure 4C, Table S5/S6). At both times points, the majority of DEGs were upregulated, with no DEG downregulated at 5 hours and only 3 genes downregulated at 24 hours. Results were subjected to gene set enrichment analysis to better understand global pathway changes due to bacterial infection (Figure 4D). There were no significantly regulated pathways at 24 hours, however at 5 hours pathways involved in coagulation, TNFα signaling, bile acid metabolism, xenobiotic metabolism and MYC targeting were all upregulated. Though they were not statistically significantly up or down-regulated statistically, GSEA analysis also indicated changes to inflammatory response, interferon alpha and gamma, TGFβ, DNA repair, complement, peroxisome, apoptosis, and angiogenesis pathways. These changes are all consistent with an inflammatory response. Though care must be taken not to over-interpret these data as they only represent the murine response to infection by cells found in the dermis and epidermis, they are overall consistent with the early genetic markers of the inflammatory response we observed in our more longitudinal models described below.

**Figure 4:**
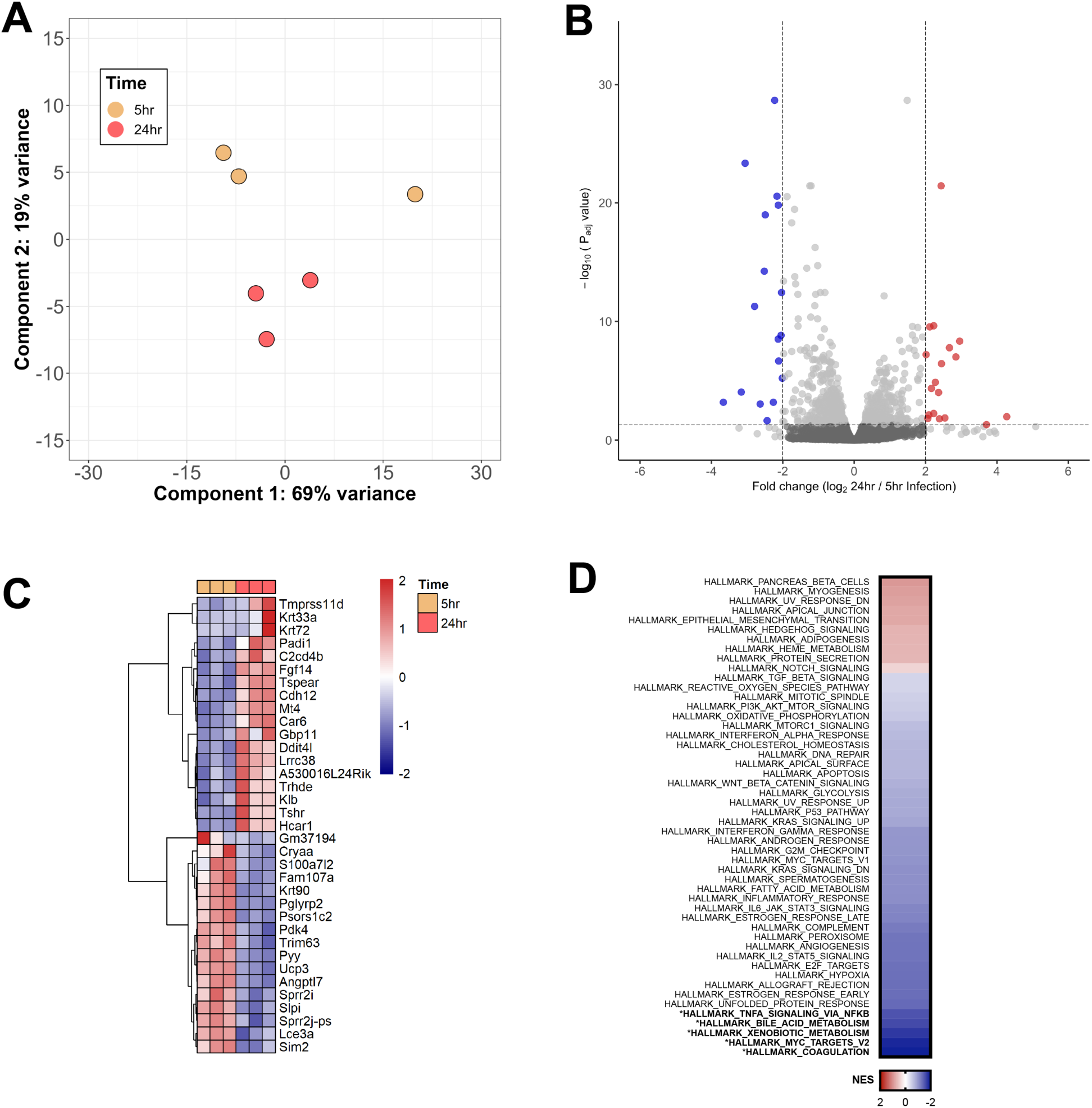
Mouse transcriptional response after 5- or 24-hours post *S. pyogenes* NZ131 skin infection. **A.** Principal-component analysis of Mouse skin infection. **B.** Differential gene expression with significant genes (Red, log_2_FC <–2 and FDR <0.05) associated with 5- or 24-hour infection (Blue, log2FC >2 and FDR <0.05). **C.** Heatmap of the top differentially expressed mouse genes which are normalized counts are transformed with Z-score method. **D.** Gene set enrichment analysis (GSEA) was performed on the fold change of 5-hour and 24-hour skin infections. Blue indicates gene sets enriched at 5-hour skin infections while red indicates gene sets enriched at 24 hour infections. Bold text indicates significance.

### An intact Rgg2/3 quorum sensing pathway is necessary to maintain NZ131 infection in a murine skin model

To provide *in vivo* validation of the results suggested by our RNAseq data, we compared infection by *S. pyogenes* with and without active Rgg2/Rgg3 QS in a modified skin inoculation model. In this model, a Tegaderm occlusive dressing was used to secure the band-aid against the murine skin for the duration of the experiment. Prior experiments demonstrated that such an approach allowed for a detectable bacterial burden for greater than 7 days (data not shown). Wild type QS-competent NZ131 were compared to a constitutively active QS Δ*rgg3* mutant, and an isogenic Δ*rgg3 shp2_GGG_shp3_GGG_* QS-null mutant. In laboratory conditions, WT NZ131 will not activate the Rgg2/Rgg3 system without stimulus (either synthetic peptides or environmental conditions, discussed below), while the Δ*rgg3* strain, lacking the negative regulator of QS, will auto-induce in a chemically defined medium [25]. Mutation of the start codon of the QS pheromones *shp2* and *shp3* renders an Δ*rgg3* strain incapable of producing the QS pheromones and thus is unable to sustain activation of the QS-associated operons [26, 27].

Compared to mice infected with QS-active strains (WT and Δ*rgg3*), mice colonized by the QS-null mutant (Δ*rgg3* shp2_GGG_shp3_GGG_) lost significantly less weight over the course of the experiment (Figure 5A). All mice demonstrated approximately 10-15% weight loss on day one. Mice inoculated with functional QS strains continued to lose weight throughout the experiment, and by day 10, these mice had lost a statistically significant greater amount of weight than those colonized by the QS-null strain. 2 mice from the WT group and 5 mice from the Δ*rgg3* group needed to be sacrificed prior to day 10 due to greater than 25% weight loss. A similar albeit delayed trend bore out regarding bacterial burden (Figure 5B). During days 1-7 of the experiment, approximately 10^7^ CFU/swab recovered in all groups. However, by day 10, the burden of QS- active (WT and Δ*rgg3*) had climbed to approximately 5×10^7^ CFU/swab, a significantly greater burden than found on mice inoculated by the QS-null strain which had declined to 8×10^6^ CFU/swab on day 10. These results demonstrate that an active Rgg2/3 QS pathway yields a survival benefit for NZ131 in the context of murine skin colonization.

**Figure 5:**
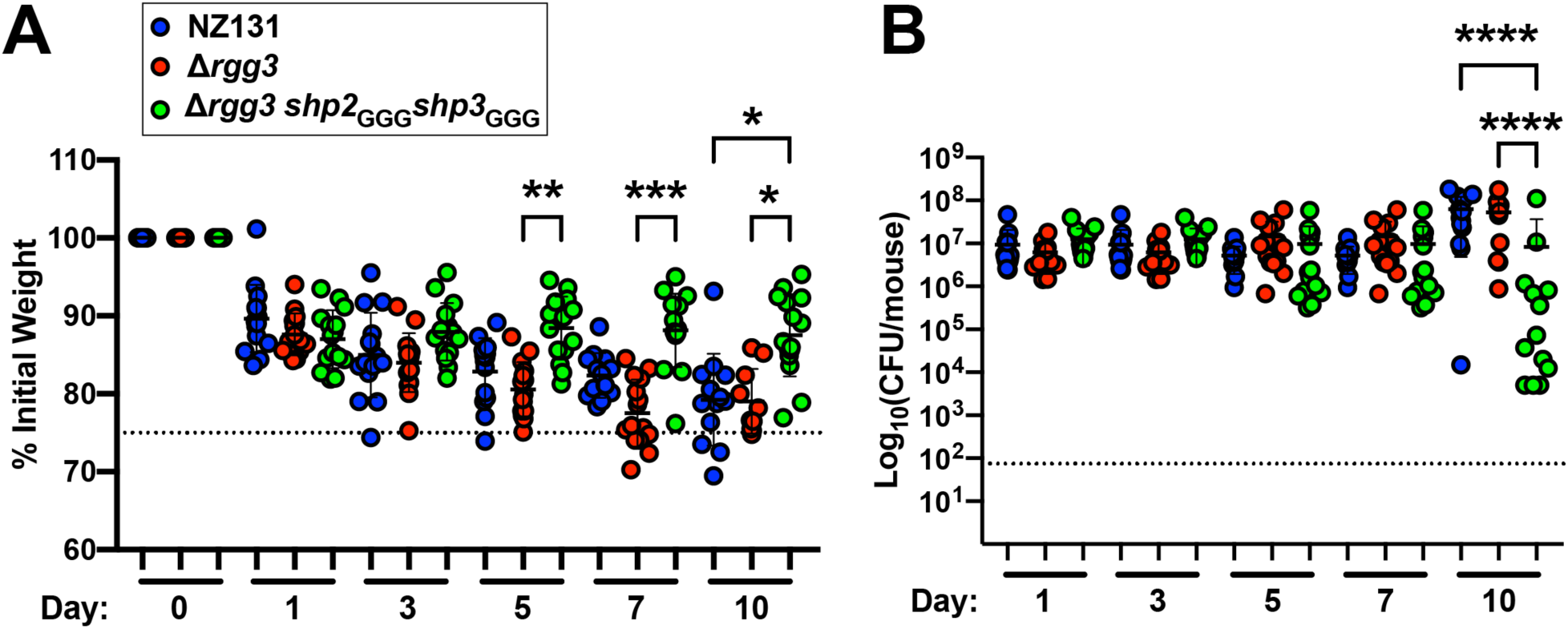
**A.** Mouse weight relative to initial weight. Mice were weighed at indicated time points, with percent weight loss calculated versus initial. Dashed line indicated 75% threshold for morbidity. **B.** CFU per mouse at indicated time points. Dashed line indicates minimum threshold of detection. For both figures, errors bars indicated mean with standard error of measurement. 15 mice per group. * P<0.05 ** P<0.005, *** P<0.0005 **** P<0.0001.

Clinical examination of murine skin throughout the experiment corresponded with the weight loss and bacterial burden data (Figure 6). On day 1, all mice (including PBS controls) were noted to have erythema along the borders of the band aid pad (Figure 6, row 1). On PBS control mice, this erythema resolved by day 3; however, mice colonized with either QS-null or QS- competent *S. pyogenes* developed diffuse erythema across the area of inoculation (Figure 6, row 2). By day 5, mice infected with QS-active strains began to show loss of epithelial barrier integrity, with lesions taking on a moist appearance (Figure 6, Row 3). These results stand in contrast to mice inoculated with the QS-null strain, who by day 5 developed crusted lesions, with clearing of the central erythema and ongoing healing continuing through day 10. Mice inoculated with QS- competent strains developed signs of worsening skin breakdown by day 10, with areas of patchy ulceration developing and a thick fibrinous layer developing over much of the inoculated area (Figure 6, row 4).

**Figure 6:**
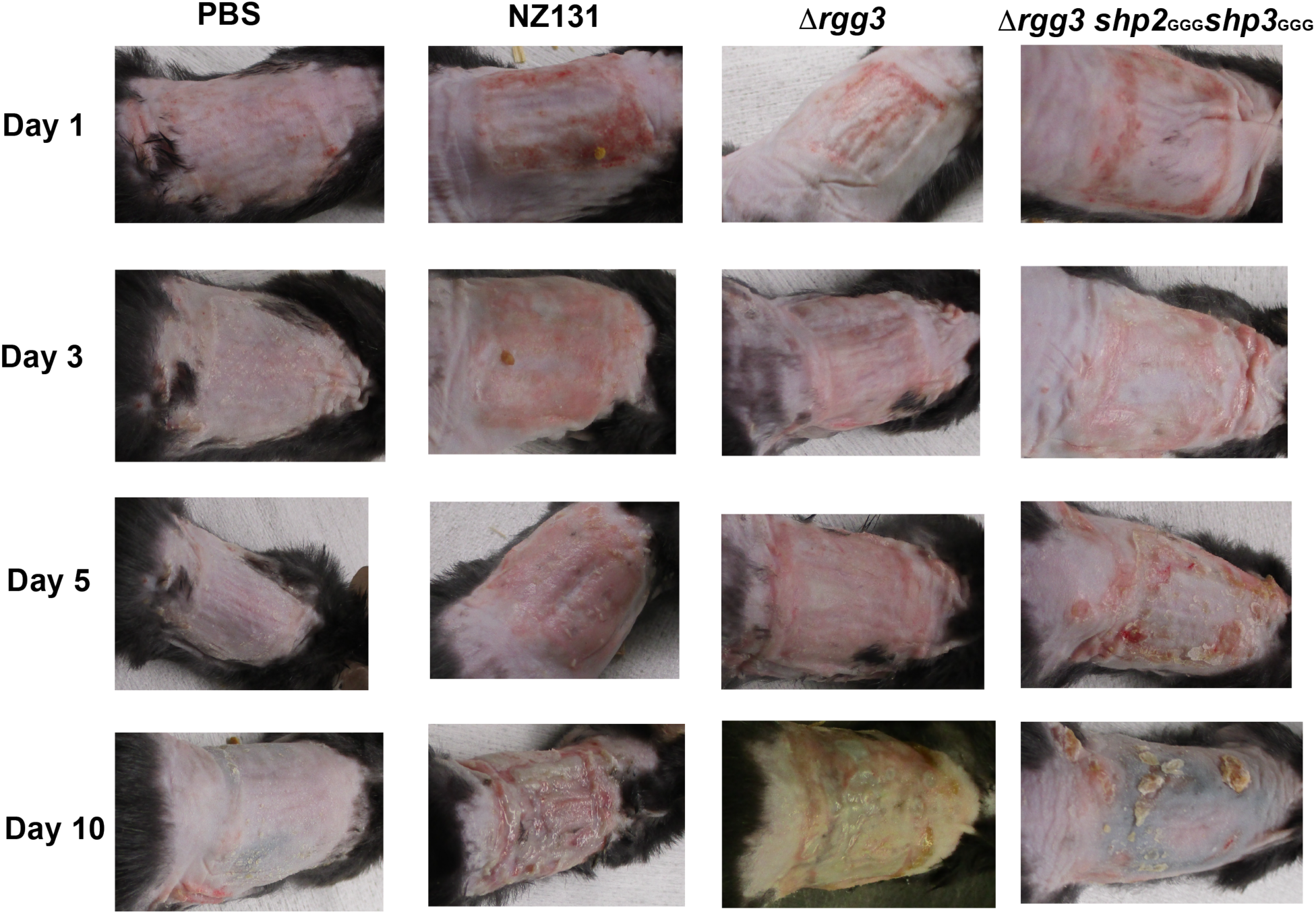
Representative photographs of murine skin colonization across time. All mice colonized with *S. pyogenes* NZ131 and isogenic mutants demonstrated superficial erythema on day 1 of the experiment. Over time, mice infected with NZ131 capable of active QS demonstrated ongoing skin breakdown, with serous discharge apparent. In comparison, mice infected with NZ131 incapable of QS initially demonstrated limited skin breakdown (Day 5), but with marked resolution by day 10.

Histology prepared from biopsies collected at day 10 is consistent with gross examination (Figure 7). Hematoxylin and eosin-stained slides show loss of the epithelial layer in mice infected by WT NZ131 and Δ*rgg3* bacterium (Figure 7, row 1). Where the epithelium remains, there is marked acanthosis. Higher power images demonstrate marked hypercellularity of mice infected by QS-active strains as compared to those infected by the QS-null strain (Figure 7, row 1 insets). Staining against myeloperoxidase (MPO, indicative of neutrophil activation) is further supportive of an inflammatory response, with strong MPO positivity noted in the subcutaneous connective tissue of mice infected by QS-active bacteria (Figure 7, row 2). Similarly, F4/80 staining against macrophages demonstrates a similar pattern (Figure 7, row 3). These microscopic results suggest a sustained or uncontrolled pro-inflammatory response by the murine epithelium when inoculated with QS-active NZ131, consistent with the changes we observed in our RNAseq GSEA results as well as providing cellular level evidence for the gross changes noted during the experiment.

**Figure 7:**
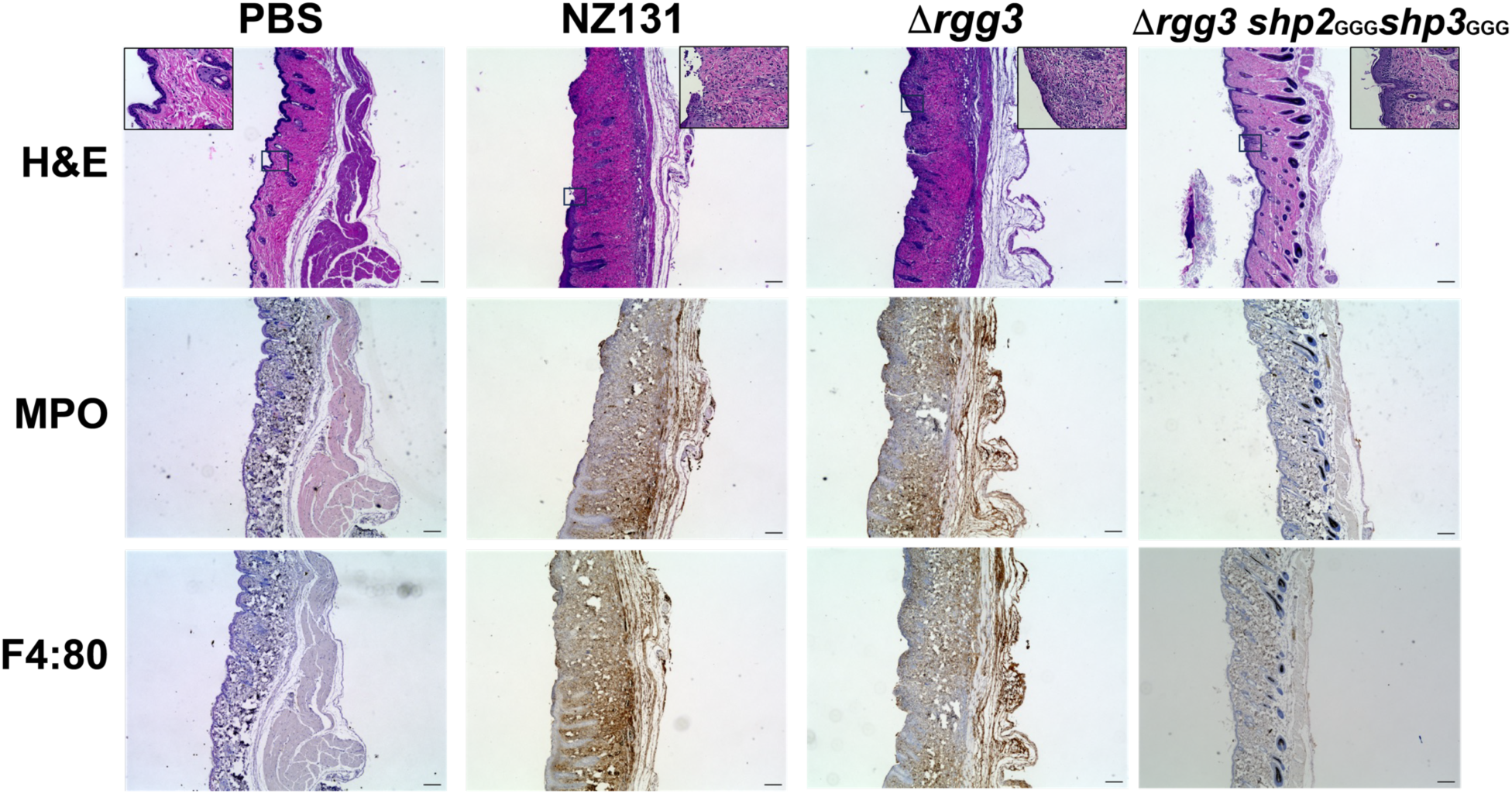
Representative microscopic images from skin biopsies taken on day 10 of longitudinal infection experiments. Samples were fixed in paraformaldehyde prior to sectioning and staining. Large mages taken at 4x fold magnification, scale bars 200 um. Boxes indicate source of inset images. Inset images taken at 40x magnification, scale bars 50 um. Images were adjusted in Image J with ‘automatic’ window and level correction for clarity.

## Discussion

*S. pyogenes* remains an enigmatic pathobiont. As a human-restricted organism, it has exquisitely evolved virulence factors and control mechanisms to best survive in its sole host. Despite years of inquiry, little is known about the early stages of bacteria-host interactions, when *S. pyogenes* is likely interacting with intact barrier surfaces (e.g. pharyngeal mucosa or integument). We set out to interrogate both host and bacterial determinants of this early yet essential step in infection. In this study, we have demonstrated that *S. pyogenes* NZ131 can transiently infect the intact murine epithelium at densities which are sufficient to isolate RNA to perform dual RNAseq, allowing us to assess both bacterial and murine transcriptomes during this interaction. Our transcriptomic data suggest that Rgg2/Rgg3 QS are up-regulated during skin infection. We have validated our RNAseq dataset with *in vivo* imaging demonstrating that murine skin infection leads to activation of the Rgg2/3 QS system at the transcriptional level. Phenotypically, infection with NZ131 capable of active Rgg2/Rgg3 QS led to more weight loss and increased bacterial burdens in an extended intact skin infection model (Figure 8). Histology derived from these experiments demonstrate an increased inflammatory response by the murine epithelium, as evidenced by mixed areas of acanthotic change and epithelial loss noted on H&E staining, as well as increased neutrophil and macrophage recruitment as evidenced by increased MPO and F4.80 staining.

**Figure 8:**
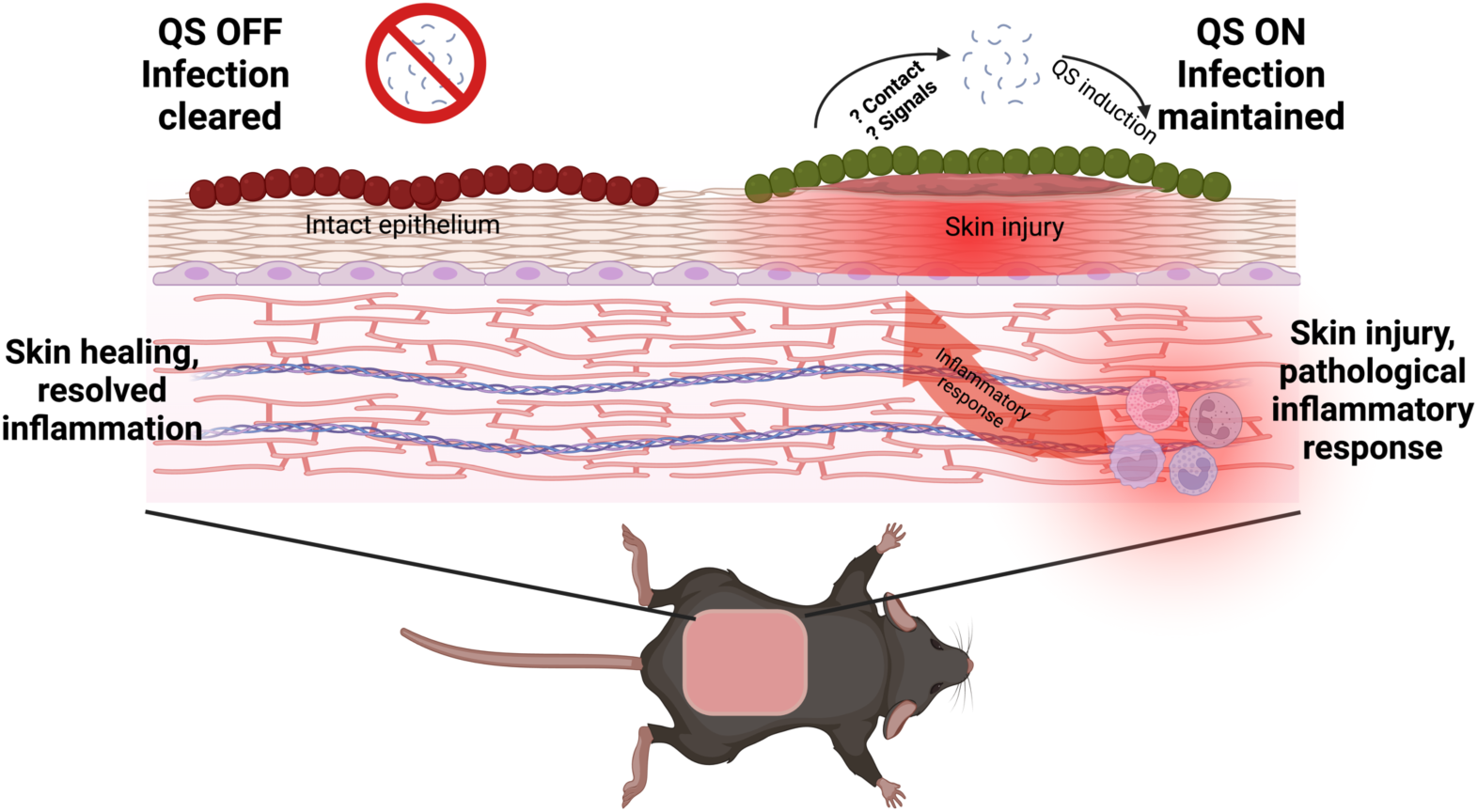
Superficial infection by *S. pyogenes* NZ131 capable of Rgg2/Rgg3 QS leads to maintenances of infection, with resulting increased skin injury, thought to be due to inflammatory infiltrate. In contrast, superficial infection by bacteria with abrogated Rgg2/Rgg3 QS are more readily cleared with resulting healing of epidermis. Created with BioRender.com.

The important connection between *S. pyogenes* skin colonization, infection and non- suppurative sequala has long been recognized. Dating back to the 1950s, the Red Lake studies monitored children and their families on the Red Lake Native American Reservation. Initially, this population was noted to have a high incidence of glomerulonephritis, which was eventually identified to be due to endemic streptococcal infections [34]. Longitudinal studies of the same population utilizing serial clinical examination, bloodwork and microbiological studies concluded that skin carriage was “a primary predisposing factor to pyoderma” [35], linking carriage to infection. Further studies concluded that *S. pyogenes* serotypes demonstrate tropism for either skin or respiratory infections, with some strains able to infect both sites [36]. This observation has been extended by the Bessen Laboratory, who demonstrated that the M protein is critical for defining tissue tropism [37]. The findings of the Red Lake Studies have been generalized in other populations and locals. A 1985 study by Maddox, Ware and Dillon conducted in a rural Alabama daycare found that 81% of their study population carried *S. pyogenes* on their skin [38]. More contemporarily, Bowen *et al*. found a close association between *S. pyogenes* and *S. aureus* nasal carriage and impetigo amongst remote Indigenous children in Australia [7] and estimate that at any one time 162 million children worldwide suffer for impetigo [39]. Superficial infections skin infections have clinically relevant long-term consequences for morbidity and mortality, with rheumatic heart disease estimated to cause 305,000 annual deaths worldwide [40, 41] and PSGN thought to be a major risk factor in the development of chronic kidney disease [1, 2, 42]. Years of studies demonstrate that *S. pyogenes* can be endemically carried on the skin, with carriage contributing to both superficial infection and high rates of non-suppurative sequelae.

To our knowledge, ours is the first study to demonstrate the roll of Rgg2/Rgg3 QS in skin infection using an unbiased RNAseq approach. The Rgg2/Rgg3 QS pathway has proven to be an enigmatic operon in *S. pyogenes*. Though this pathway is conserved amongst all sequenced isolates of *S. pyogenes*, it has been challenging to determine *in vivo* phenotypes associated with operon activation. Prior studies defined new *in vitro* functions for this QS pathway, including increased lysozyme resistance, decreased *slo* expression, and impaired macrophage activation [32]. It was also determined that the Rgg2/3 pathway is regulated indirectly by *covRS* via the degradation of the Shp pheromones by the endopeptidase PepO [18, 31–33]. Gogos *et al.* contributed *in vivo* evidence of Rgg2/3 function by demonstrating that active Rgg2/Rgg3 QS led to greater bacterial burden in a murine NALT model, with an associated increase in mortality [33]. Concordant to our study, Gogos *et al.* also found *in vivo* QS activation in the NALT with IVIS, though they proposed their studies as one of colonization. Collectively, these results have led to the hypothesis that the Rgg2/Rgg3 pathway may play a role in early infection and immune evasion.

Though we initially envisioned our model as one of carriage, both the gross and microscopic images from our longitudinal infection model would suggests that Rgg2/Rgg3 QS activation leads to a more morbid phenotype than QS-null infection. From our current studies, we cannot discern if this is due to an increase in virulence or simply as a side effect of persistence. As discussed above, genes associated with virulence did not show a consistent trend in upregulation during skin infection, nor do other genes associated with the master regulator of virulence *covRS*. Though it remains a possibility the transcriptional landscape changes beyond 24 hours post infection, based on these data we favor the hypothesis that Rgg2/Rgg3 activation provides a more hospitable environment for NZ131 infection. Either by improving bacterial fitness in this challenging environment or altering host immune response, QS-active bacteria are unable to be effectively cleared. Additional work is needed to better understand the mechanism of this phenotype and dissect increased virulence from persistence.

The mechanism by which murine skin colonization drives Rgg2/3 QS activation remains an area of active inquiry. Prior work has demonstrated that high-mannose low-glucose conditions can induce Rgg2/3 QS via Mga. Phosphotransferase (PTS) signaling under mannose conditions alters the Mga phosphorylation state, enhancing Mga’s ability to repress transcription of Rgg3 [43]. This mechanism presumably allows NZ131 to fine-tune its response to host-environment carbon sources. The RNAseq data presented herein showed that *rgg3*, the negative regulator of QS, was approximately 5-fold downregulated at both 5 and 24 hours post murine infection and we therefor speculate this may represent Mga-mediated *rgg3* repression as described by Woo *et al* [43]. The complete Mga regulon in NZ131 has not been fully characterized; however, a core regulon for *S. pyogenes* has been described. Such genes include autoregulation of *mga,* upregulation of *emm, scpA, sclA, salA* and *fba* [44–46]. In our RNAseq data, these genes are not consistently upregulated as would be predicted via direct *mga* activation. We also considered the metal-response regulator MtsR could be mediating QS induction, as free iron is tightly regulated in both humans and mice. In culture conditions where iron is replete, MtsR binds at the *shp3* promoter and blocks QS induction. MtsR regulates other *S. pyogenes* genes in response to iron availability, repressing *nrdF2* and *Ska*, while upregulating *pyrFE* when in iron-replete conditions [47, 48]. Although we observed upregulation of *nrdF2* in our RNAseq dataset, no change were seen for *ska* or *pyrFE*. Of note, there is significant overlap between the Mga and MtsR regulons, thus our data could represent a combination of effector functions. Additional experiments are needed to elucidate the mechanism by which skin contact induces Rgg2/Rgg3 QS.

Additional studies are needed to determine if the findings described herein extend to human epithelial interactions. Though murine epithelium is a useful experimental substrate, there are several key differences between the human and murine integument, including the conspicuous absence of eccrine sweat glands and neutrophil-derived defensins in the mouse. Furthermore, human skin is 5-10 cells thick with attachment to underlying tissue, while murine skin is only 2-3 cells thick with increased permeability and decreased barrier functionality, as well as no direct attachment to underlying tissue [49, 50]. As an obligate human pathobiont, *S. pyogenes* has co-evolved to bind human adhesins while mitigating the human immune response. Despite the differences between human and murine epithelium, neither a recent myositis model in non-human primates nor a vaginal murine carriage model using *S. pyogenes* found Rgg2/Rgg3 activation in their transcriptomic studies, suggesting that the interaction between *S. pyogenes* and the skin may uniquely initiate Rgg2/Rgg3 activation [10, 23].

Though the findings of this study that Rgg2/Rgg3 activity led to prolonged duration of infection and increased bacterial burden seem discordant with the previously proposed role for Rgg2/Rgg3 in colonization [33], we propose that improved immune evasion by QS-active *S. pyogenes* allows for the development of a greater bacterial burden in this model, thus explaining the increased weight loss and bacterial burden noted in mice colonized with QS-active bacteria. Additional work is needed to elucidate the mechanism by which the Rgg2/Rgg3 regulon enhances maintenance of infection, especially regarding which genes under Rgg2/Rgg3 regulation are necessary and sufficient for this phenotype. A more complete understanding of this pathway may offer novel therapeutic approaches to preventing *S. pyogenes* colonization or infection using a quorum sensing inhibition approach, and indeed QS inhibitors have already been identified [51]. Given the streptococcal disease burden demonstrated in both historical studies (such as the Red Lake studies) as well as more recent studies of Aboriginal populations in Australia and New Zealand, and the ongoing impact of both invasive *S. pyogenes* and non- suppurative sequalae, there is a clear need for further work to elucidate the mechanisms by which *S. pyogenes* colonizes and infects healthy skin.

## Supplementary Tables

S1: Complete RNAseq results from 5-hour samples, aligned to the NZ131 genome NC_011375.1. Name refers to current nomenclature per NCBI, while old_locus refers to historical annotation.

S2: Complete RNAseq results from 24-hour samples, aligned to the NZ131 genome NC_011375.1. Name refers to current nomenclature per NCBI, while old_locus refers to historical annotation.

S3: RNAseq results from 5-hour samples, filtered for genes with a log_2_ FC >|2| and padj<0.05 S4: RNAseq results from 24-hour samples, filtered for genes with a log_2_ FC >|2| and padj<0.05 S5: Complete RNAseq results from 5-hour samples, aligned to the *M. musculus* genome mm1. S6: Complete RNAseq results from 24-hour samples, aligned to the *M. musculus* genome mm1.

## Acknowledgments

The authors thank the members of the Horswill and Doran Labs for their assistance in the design of these studies and preparation of this manuscript. The Federle Lab provided the strains used in these experiments, as well as a fundamental understanding of the Rgg2Rgg3 QS pathway. Laura Hoaglin of the Dermatohistology Core was essential for slide preparation and staining. Grant support came from NIH NIAID grants AI153185 and AI162964 to Dr. Horswill and salary support to Dr. Wilkening came from the Section of Pediatric Critical Care Medicine in the Department of Pediatrics at the University of Colorado School of Medicine.

## References

1. Cunningham, M.W., Pathogenesis of group A streptococcal infections. Clin Microbiol Rev, 2000. 13(3): p. 470–511.

2. Carapetis, J.R., et al., The global burden of group A streptococcal diseases. Lancet Infect Dis, 2005. 5(11): p. 685–94.

3. Bessen, D.E., Population biology of the human restricted pathogen, Streptococcus pyogenes. Infect Genet Evol, 2009. 9(4): p. 581–93.

4. DeMuri, G.P. and E.R. Wald, The Group A Streptococcal Carrier State Reviewed: Still an Enigma. J Pediatric Infect Dis Soc, 2014. 3(4): p. 336–42.

5. Byrd, A.L., Y. Belkaid, and J.A. Segre, The human skin microbiome. Nat Rev Microbiol, 2018. 16(3): p. 143–155.

6. Luna, P.C., Skin Microbiome as Years Go By. Am J Clin Dermatol, 2020. 21(Suppl 1): p. 12–17.

7. Bowen, A.C., et al., The microbiology of impetigo in indigenous children: associations between Streptococcus pyogenes, Staphylococcus aureus, scabies, and nasal carriage. BMC infectious diseases, 2014. 14(1): p. 727–727.

8. Swe, P.M., et al., Complement inhibition by Sarcoptes scabiei protects Streptococcus pyogenes - An in vitro study to unravel the molecular mechanisms behind the poorly understood predilection of S. pyogenes to infect mite-induced skin lesions. PLoS Negl Trop Dis, 2017. 11(3): p. e0005437.

9. Hirose, Y., et al., Streptococcus pyogenes Transcriptome Changes in the Inflammatory Environment of Necrotizing Fasciitis. Appl Environ Microbiol, 2019. 85(21).

10. Zhu, L., et al., Gene fitness landscape of group A streptococcus during necrotizing myositis. J Clin Invest, 2019. 129(2): p. 887–901.

11. Team, R.C., R: A language and environment for statistical computing. 2022, R Foundation for Statistical Computing: Vienna, Austria

12. Love, M.I., W. Huber, and S. Anders, Moderated estimation of fold change and dispersion for RNA-seq data with DESeq2. Genome Biol, 2014. 15(12): p. 550.

13. Korotkevich, G., et al., Fast gene set enrichment analysis. bioRxiv, 2021: p. 060012.

14. Liberzon, A., et al., Molecular signatures database (MSigDB) 3.0. Bioinformatics, 2011. 27(12): p. 1739–40.

15. Dolgalev, I., MSigDB Gene Sets for Multiple Organisms in a Tidy Data Format. 2022

16. Kanehisa, M. and S. Goto, KEGG: kyoto encyclopedia of genes and genomes. Nucleic Acids Res, 2000. 28(1): p. 27–30.

17. Kanehisa, M., et al., KEGG for taxonomy-based analysis of pathways and genomes. Nucleic Acids Res, 2023. 51(D1): p. D587–D592.

18. Wilkening, R.V., J.C. Chang, and M.J. Federle, PepO, a CovRS-controlled endopeptidase, disrupts Streptococcus pyogenes quorum sensing. Mol Microbiol, 2016. 99(1): p. 71–87.

19. McShan, W.M., et al., Genome sequence of a nephritogenic and highly transformable M49 strain of Streptococcus pyogenes. J Bacteriol, 2008. 190(23): p. 7773–85.

20. Bessen, D.E., et al., Genetic correlates of throat and skin isolates of group A streptococci. J Infect Dis, 1996. 173(4): p. 896–900.

21. Bessen, D.E., et al., Whole-genome association study on tissue tropism phenotypes in group A Streptococcus. J Bacteriol, 2011. 193(23): p. 6651–63.

22. Kanehisa, M., Toward understanding the origin and evolution of cellular organisms. Protein Sci, 2019. 28(11): p. 1947–1951.

23. Cook, L.C.C., et al., Transcriptomic Analysis of Streptococcus pyogenes Colonizing the Vaginal Mucosa Identifies hupY, an MtsR-Regulated Adhesin Involved in Heme Utilization. mBio, 2019. 10(3).

24. Finn, M.B., et al., Identification of group a streptococcus genes directly regulated by csrrs and novel intermediate regulators. mBio, 2021. 12(4).

25. Chang, J.C., et al., Two group A streptococcal peptide pheromones act through opposing Rgg regulators to control biofilm development. PLoS Pathog, 2011. 7(8): p. e1002190.

26. LaSarre, B., J.C. Chang, and M.J. Federle, Redundant group a streptococcus signaling peptides exhibit unique activation potentials. J Bacteriol, 2013. 195(18): p. 4310–8.

27. Lasarre, B., C. Aggarwal, and M.J. Federle, Antagonistic Rgg regulators mediate quorum sensing via competitive DNA binding in Streptococcus pyogenes. mBio, 2013. 3(6).

28. Jimenez, J.C. and M.J. Federle, Quorum sensing in group A Streptococcus. Front Cell Infect Microbiol, 2014. 4: p. 127.

29. Aggarwal, C., et al., Multiple length peptide-pheromone variants produced by Streptococcus pyogenes directly bind Rgg proteins to confer transcriptional regulation. J Biol Chem, 2014. 289(32): p. 22427–36.

30. Wilkening, R.V. and M.J. Federle, Evolutionary Constraints Shaping Streptococcus pyogenes-Host Interactions. Trends Microbiol, 2017. 25(7): p. 562–572.

31. Chang, J.C., et al., Quorum Sensing Regulation of a Major Facilitator Superfamily Transporter Affects Multiple Streptococcal Virulence Factors. J Bacteriol, 2022. 204(9): p. e0017622.

32. Rahbari, K.M., J.C. Chang, and M.J. Federle, A Streptococcus Quorum Sensing System Enables Suppression of Innate Immunity. mBio, 2021. 12(3).

33. Gogos, A. and M.J. Federle, Colonization of the Murine Oropharynx by Streptococcus pyogenes Is Governed by the Rgg2/3 Quorum Sensing System. Infect Immun, 2020. 88(10).

34. Reinstein, C.R., Epidemic nephritis at Red Lake, Minnesota. J Pediatr, 1955. 47(1): p. 25–34.

35. Dudding, B.A., et al., The role of normal skin in the spread of streptococcal pyoderma. J Hyg (Lond), 1970. 68(1): p. 19–28.

36. Bascom, A., et al., THE DYNAMICS OF STREPTOCOCCAL INFECTIONS IN A DEFINED POPULATION OF CHILDREN: SEROTYPES ASSOCIATED WITH SKIN AND RESPIRATORY INFECTIONS. American Journal of Epidemiology, 1976. 104(6): p. 652–666.

37. Bessen, D.E., Tissue tropisms in group A Streptococcus: What virulence factors distinguish pharyngitis from impetigo strains? 2016, Lippincott Williams and Wilkins. p. 295–303.

38. Maddox, J.S., J.C. Ware, and H.C. Dillon, Jr., The natural history of streptococcal skin infection: prevention with topical antibiotics. J Am Acad Dermatol, 1985. 13(2 Pt 1): p. 207–12.

39. Bowen, A.C., et al., The Global Epidemiology of Impetigo: A Systematic Review of the Population Prevalence of Impetigo and Pyoderma. PLoS One, 2015. 10(8): p. e0136789.

40. de Loizaga, S.R., et al., Rheumatic Heart Disease in the United States: Forgotten But Not Gone: Results of a 10 Year Multicenter Review. J Am Heart Assoc, 2021. 10(16): p. e020992.

41. de Loizaga, S.R. and A.Z. Beaton, Rheumatic Fever and Rheumatic Heart Disease in the United States. Pediatr Ann, 2021. 50(3): p. e98–e104.

42. Hoy, W.E., et al., Post-streptococcal glomerulonephritis is a strong risk factor for chronic kidney disease in later life. Kidney Int, 2012. 81(10): p. 1026–1032.

43. Woo, J.K.K., K.S. McIver, and M.J. Federle, Carbon catabolite repression on the Rgg2/3 quorum sensing system in Streptococcus pyogenes is mediated by PTS(Man) and Mga. Mol Microbiol, 2022. 117(2): p. 525–538.

44. Flores, A.R., et al., Natural variation in the promoter of the gene encoding the Mga regulator alters host-pathogen interactions in group a Streptococcus carrier strains. Infect. Immun., 2013. 81(11): p. 4128–4138.

45. Hondorp, E.R. and K.S. McIver, The Mga virulence regulon: infection where the grass is greener. Mol Microbiol, 2007. 66(5): p. 1056–65.

46. Ribardo, D.A. and K.S. McIver, Defining the Mga regulon: Comparative transcriptome analysis reveals both direct and indirect regulation by Mga in the group A streptococcus. Mol Microbiol, 2006. 62(2): p. 491–508.

47. Toukoki, C., et al., MtsR is a dual regulator that controls virulence genes and metabolic functions in addition to metal homeostasis in the group A streptococcus. Mol Microbiol, 2010. 76(4): p. 971–89.

48. Bates, C.S., et al., Characterization of MtsR, a new metal regulator in group A streptococcus, involved in iron acquisition and virulence. Infect Immun, 2005. 73(9): p. 5743–53.

49. Zomer, H.D. and A.G. Trentin, Skin wound healing in humans and mice: Challenges in translational research. J Dermatol Sci, 2018. 90(1): p. 3–12.

50. Medetgul-Ernar, K. and M.M. Davis, Standing on the shoulders of mice. Immunity, 2022. 55(8): p. 1343–1353.

51. Aggarwal, C., et al., Identification of Quorum-Sensing Inhibitors Disrupting Signaling between Rgg and Short Hydrophobic Peptides in Streptococci. mBio, 2015. 6(3): p. e00393–15.

52. Simon, D. and J.J. Ferretti, Electrotransformation of Streptococcus pyogenes with plasmid and linear DNA. FEMS microbiology letters, 1991. 66(2): p. 219–224.

53. Cook, L.C., B. LaSarre, and M.J. Federle, Interspecies communication among commensal and pathogenic streptococci. mBio, 2013. 4(4): p. e00382--13--e00382--13.

